# Quantify nanoscopic alterations associated with mitochondrial dysfunctions using three-dimensional single-molecule localization microscopy

**DOI:** 10.1101/2023.05.10.540227

**Authors:** Benjamin Brenner, Yang Zhang, Junghun Kweon, Raymond Fang, Nader Sheibani, Sarah X. Zhang, Cheng Sun, Hao F. Zhang

**Affiliations:** Department of Biomedical Engineering, Northwestern University, Evanston, IL; Department of Ophthalmology and Vision Sciences, University of Wisconsin, Madison, WI; Department of Ophthalmology, University at Buffalo, Buffalo, NY; Department of Mechanical Engineering, Northwestern University, Evanston, IL; Currently with Program of Polymer and Color Chemistry, Department of Textile Engineering, Chemistry and Science, North Carolina State University, Raleigh, NC

## Abstract

The morphology of mitochondria provides insights into their functions. Three-dimensional (3D) single-molecule localization microscopy (SMLM) uniquely enables the analysis of mitochondrial morphological features individually, thanks to its high resolution. However, nearly all reported studies of mitochondrial morphology have been qualitative and without statistical analysis. We report a quantitative method to extract mitochondrial morphological metrics, including volume, aspect ratio, and local protein density, from 3D SMLM images, with single-mitochondrion sensitivity. We validated our approach using simulated ground-truth SMLM images of mitochondria. We further tested our morphological analysis on mitochondria that have been altered functionally and morphologically in controlled manners. This work sets the stage to analyze mitochondrial morphological alterations associated with disease progression quantitatively.

## INTRODUCTION

Fluorescence microscopy is widely used in cell and molecular biology to selectively label and image cellular organelles and proteins of interest. Several fluorescence imaging studies have linked cellular function with spatial information about their organelles [1-3]. Mitochondria are of particular interest as mitochondrial dysfunction has been linked to numerous pathologies, including mitochondrial diseases [4], cancer [5, 6], diabetes [7, 8], diabetic complications [9-13], Alzheimer’s disease [14, 15], and Parkinson’s disease [16-19]. It has been long postulated that a significant correlation exists between mitochondrial structure and functions [20, 21].

Morphological imaging of mitochondria using fluorescence microscopy has been widely used to draw important conclusions about mitochondrial functions. Olichon *et al.* used confocal microscopy images of OPA1 downregulated cells showing mitochondrial network fragmentation to support their conclusion that OPA1 plays a role in organizing the mitochondrial inner membrane [22]. Visser *et al.* used confocal microscopy to examine the morphological effects of growth conditions in fungal mitochondria [23]. Stoianovski *et al.* noted that the extent of mitochondrial fission was related to human fis1 expression in COS-7 cells, indicating that it plays a role in mitochondrial fission [24].

Among all mitochondria’s morphological alterations, the most metabolically relevant events are fusion and fission [21]. In mitochondrial fusion, cytoskeleton-dependent transport proteins bring two discrete mitochondria together. Next, the inner and outer membranes are deformed and fused by the controlled placement of hydrophobic domains on the inner and outer membranes and various fusion proteins that draw the lipid bilayers together. In mitochondrial fission, the inner and outer mitochondrial membranes of a single, continuous mitochondrion are pinched off, forming two separate mitochondria. Mitochondrial fusion promotes the exchange of complementary genetic products, minimizing stress-related mitochondrial damage. In contrast, mitochondrial fission ensures the distribution of mitochondria throughout the cell, is an essential step in cell apoptosis, and promotes mitophagy to degrade damaged or ineffective mitochondria. Researchers believe that elongated mitochondria from fusion have a higher metabolic capacity than fragmented mitochondria [25]; however, total inhibition of fission also results in lower metabolic capacity due to the accumulation of ineffective mitochondria from a lack of mitophagy [21].

Several groups developed tools, such as MitoSegNet [26], Mitohacker [27], and mitometer [28], to assess mitochondria imaged by wide-field or confocal microscopy with diffraction-limited spatial resolution rather than super-resolution imaging. As a result, these tools generated anomalous results when applied to pointillist Single-molecule localization microscopy (SMLM) images. They also cannot perform analyses only feasible at the super-resolution scale, such as identifying the interface between two individual mitochondria or precisely measuring individual mitochondrial shape and volume. To date, most biological studies that conclude morphological changes in mitochondria images rely on a qualitative evaluation of mitochondrial shape [29-32].

SMLM [33, 34] is a fluorescence microscopy technique that achieves super-resolution at a spatial precision of ∼15 nm using samples labeled with stochastically blinking fluorophores. In each frame of an SMLM dataset, only a sparse subset of fluorophores emits light, and each fluorescence emission yields a point spread function (PSF) on an array detector, usually approximated by a two-dimensional (2D) Gaussian function. The peak of this Gaussian function is treated as the true location of the fluorescence emitter. Maximum likelihood estimation is often used to localize each emitter by identifying the Gaussian peak with high precision [35]. The final SMLM image is a histogram consisting of all localizations of fluorophores. Through novel optical designs to manipulate the PSFs, researchers also achieved three-dimensional (3D) SMLM [36-38]. Furthermore, as SMLM images are histograms of discontinuous localizations of discrete fluorescence emissions, SMLM provides unique information about local protein distribution and density [39]. This can be particularly interesting when evaluating mitochondrial outer membrane cohesion, which correlates closely with mitochondrial health [40, 41].

As super-resolution imaging is increasingly incorporated into studies that draw functional and pathological information from intracellular structures [42-44], methodologies must quantitatively correlate pathological alterations with morphological parameters at the nanoscopic scale. In this study, we show that 3D SMLM can detect changes in mitochondrial volume and 3D morphology and alterations in mitochondrial membrane protein density. To demonstrate these abilities, we simulate 3D SMLM images of mitochondria with various morphologies and protein densities. We also show that 3D SMLM can detect drug-induced changes in mitochondrial morphology associated with membrane potential drop induced by ATP synthase inhibition and alterations in protein density associated with the early stages of apoptosis [45].

## METHODS AND MATERIALS

### SMLM imaging system

We built our 3D SMLM imaging system on an inverted microscope body (Ti-2, Nikon) with wide-field illumination. Fig. S1(A) shows the schematic for our experimental system using a cylindrical lens. Light from a CW 647-nm laser (2RU-VFL-P-1000-647-B1R, MPB Communications Inc.) first passed through a band-pass filter (FF01-642/10-25, Semrock), then was reflected by a 649-nm cutoff dichroic mirror (FF649-Di01-25x36, Semrock) and focused by an objective lens (OBJ. CFI SR HP Apochromat TIRF 100XC Nikon) on the sample. The emitted fluorescence photons passed through a 200-mm tube lens and a 647.1-nm long-pass filter (BLP01-647R-25, Semrock). The fluorescence photons then passed through two lenses (L1, 100-mm focal length; L2, 100-mm focal length) and a cylindrical lens (CL, 200-mm focal length) to be focused onto an electron-multiplying charge-coupled device (EMCCD, iXon Ultra 897).

### Axial calibration

We performed axial calibration using red fluorescent nanospheres (F8807, Thermo Fisher). We scanned the nanospheres across a 2-μm axial range with an interval of 20 nm. Then, we used maximum likelihood estimation [35] to fit an anisotropic 2D Gaussian function to the image of the nanosphere in each frame. Finally, we plotted a calibration curve of the Gaussian sigma as a function of the nanosphere’s axial position relative to the objective lens (Fig. S1B.)

### Characterization of axial and lateral resolutions

To characterize the axial resolution, we imaged nanospheres and added a neutral density filter to the excitation light path to ensure the nanospheres emitted 2,000 photons during each CCD exposure (10 ms). We adjusted the axial position of the objective lens focus from -400 nm to 400 nm with an interval of 200 nm (5 positions total) and captured 100 frames at each axial position. We then estimated the axial position of the nanosphere in each frame using the axial calibration curve and measured the standard deviation at each position. We multiplied this number by 2.355 (the ratio between the full-width-half-maxium or FWHM and the standard deviation of a Gaussian function) to get the axial resolution (Fig. S1C) [46], which averaged 80 nm in our mitochondrial imaging studies. Finally, we characterized lateral uncertainty using the method introduced by Rieger *et al.* [35], obtaining a mean value of 17 nm in cellular imaging, as shown in Fig. S1D.

### Sample preparation

We grew HeLa cells in cell culture flasks (T-75 Thermo Fisher) to 80% confluency in a DMEM medium. Then we detached them using trypsin and plated a 1:10 dilution in DMEM on an 8-well chamber (Lab-Tek 1.0 borosilicate glass, Thermo Fisher), which we let grow overnight. We fixed the cells with 10% PFA for 10 minutes at room temperature, permeabilized the cell membranes with 0.5% Triton-X-100 for 10 minutes, and blocked the cells with 2.5% goat serum for 30 minutes. Next, we prepared a 1:100 dilution of rabbit anti-TOM20 antibody (HPA011562 Millipore Sigma) [47] in 2.5% goat serum and incubated the cells on a rocker overnight at 4°C. We rinsed the cells three times with a washing buffer (0.2% BSA, 0.1% Triton X-100 in PBS) for 5 minutes at RT. We then added a 1:200 dilution of AF-647-conjugated goat anti-rabbit antibody (A-21245, Invitrogen) in blocking buffer to each well and incubated the cells for 45 minutes at room temperature. Finally, we rinsed the cells twice with washing buffer and twice with PBS for 5 minutes at room temperature.

### Image acquisition and image processing

We added a buffer made from 50 mM Tris, 10mM NaCl, 0.5 mg/mL glucose oxidase (Sigma, G2133), 2000 U/mL catalase (Sigma, C30), 10% (w/v) D-glucose, and 100 mM cysteamine to the samples to induce fluorescent blinking. During data acquisition, we replaced the buffer every two hours. Our excitation wavelength was 647 nm with an excitation power of 45 W/mm^2^. We captured 10,000 frames with 20-ms exposure time. For each field of view (FOV), we acquired two sets of frames: one with the objective focused at -250 nm and one with the objective focused at +250 nm. This acquisition strategy ensured that the axial depth of field was sufficiently large to capture all the mitochondria in each cell.

We used the ImageJ plug-in Thunderstorm [48] to localize the astigmatic PSFs, using the maximum likelihood function to fit a 2D ellipsoid Gaussian to each PSF [36]. When the cell size was larger than the FOV, we captured multiple FOVs with overlapping areas and stitched the images. To stitch two images, we first computed the 2D cross-correlation of the images, which we normalized by the total number of overlapping pixels at each coordinate. Next, we found the coordinates of the maximum of this 2D map from which we subtracted the size of the first image. We then translated the second image by the resulting value.

### Mitochondrial membrane potential (MMP) measurement

We plated 15,000 HeLa cells per well in a 96-well plate (3894, Costar) and incubated them overnight at 37°C. Next, we washed the wells with PBS, added 10 μM JC-1 (tetraethylbenzimidazolylcarbocyanine iodide), an MMP reporter dye that targets mitochondria [49] (ThermoFisher, Mitoprobe JC-1 assay kit), to the control and treated groups, and incubated them for 10 minutes at 37°C. Next, we washed the wells three times with PBS and added 100 μM FCCP (carbonyl cyanide 4-(trifluoromethoxy) phenylhydrazone) to the FCCP treated group, 10-μM staurosporine (STS) to the STS treated group, and 2.5-μM oligomycin and 1-μM antimycin (OA) [26, 61] to the OA treated group in dilution buffer (ab113850, Abcam) with 10% fetal bovine seru for 3 hours at 37° to induce fragmentation and lower MMP. Within 10 minutes, we transferred the plate to a plate reader (Cytation 5, BioTek) at 37°C. We set the plate reader to document fluorescence with an excitation wavelength of 475 ± 10 nm and an emission of 590 ± 10 nm with an interval of 2 minutes for four hours. Finally, we took the ratio of fluorescence intensities in the 590-nm channel to fluorescence intensities in the 475-nm channel, referred to as the relative fluorescent units (RFU), as a metric of MMP.

### Three-dimensional confocal microscopy

We performed 3D confocal microscopy using a Leica SP5 microscope. We excited the samples using a 633 nm excitation and captured fluorescence emissions from 650 nm to 710 nm. We used a pixel size of 116 μm and performed 3-line averaging and then 2-frame averaging. We acquired a depth range of 4 μm with an interval of 125 nm.

## RESULTS

Fig. 1 illustrates mitochondrial fragmentation, biochemical validation of JC-1 for measurement of MMP, and resolution comparison between 3D SMLM and confocal microscopy. Figs. 1A & 1B illustrate how mitochondrial fission results in a more fragmented mitochondrial morphology. Fig. 1C shows that MMPs in untreated cells remained relatively stable compared to the decay in MMP in cells treated with FCCP, the depolarization control [49], as indicated by the normalized RFU time course. In the control group, variations in the JC-1 signal with an increased MMP may be attributed to movement of the cells from a room temperature environment to the 37°C environment inside the plate reader [50], while the subsequent drop in JC-1 signal may have been a combined effect of JC-1 dissociation from the mitochondria and photobleaching. Fig. 1D is the 2D projection confocal image of an untreated HeLa cell with a color map to indicate axial position, and Fig. 1E is the magnified view of the highlighted region in Fig. 1D.

**Fig. 1.**
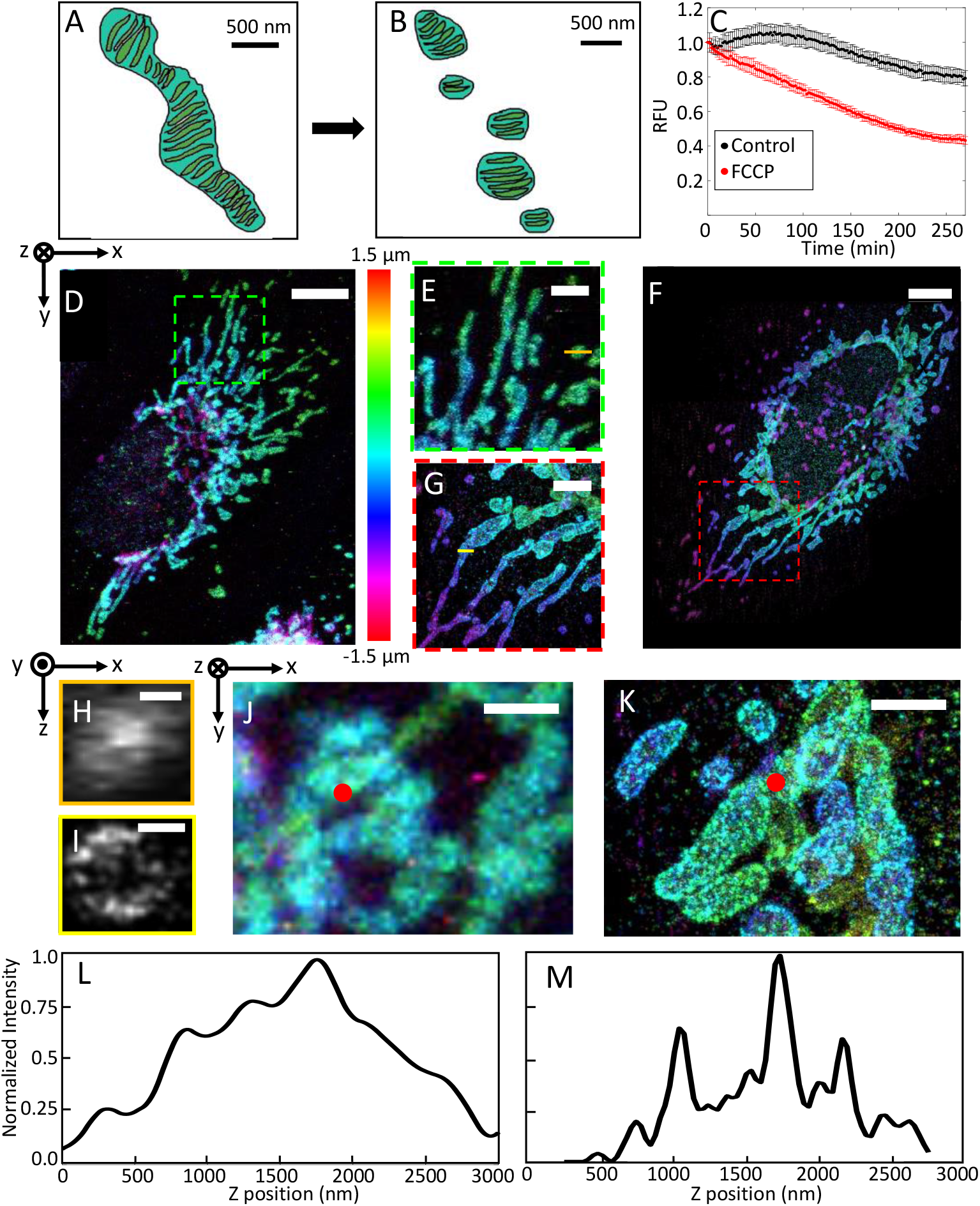
Illustration of (A) an elongated mitochondrion and (B) fragmented mitochondria. The intermembrane space is colored turquoise, and the cristae are colored green; (C) Normalized RFU of JC-1-labeling over time of untreated (black) and FCCP-treated (red) HeLa cells; (D) projection view of mitochondria imaged by 3D confocal microscopy with a color bar to indicate z-position. Scale bar: 5 μm; (E) Magnified view of the area highlighted in panel D. Scale bar: 2.5 μm; (F) projection view of mitochondria imaged by 3D SMLM. Scale bar: 5 μm; (G) Magnified view of the area highlighted in panel D. Scale bar: 2.5 μm; (H) Cross-section view of one mitochondrion imaged by confocal microscopy from the position highlighted in panel E. Scale bar: 250 nm; (I) Cross-section view of one mitochondrion imaged by SMLM from the position highlighted in panel G. Scale bar: 250 nm; (J) Projection view of densely-packed mitochondria imaged by confocal microscopy. Scale bars: 2 μm; (K) Projection view of densely-packed mitochondria imaged by SMLM. Scale bars: 2 μm; (L) Depth profile of stacked mitochondria from the position highlighted in panel J. (M) Depth profile of stacked mitochondria from the position highlighted in panel K.

Figs. 1F & 1G show the 2D projection of 3D SMLM images of an untreated HeLa cell (Fig. 1G is the magnified view of the highlighted region in Fig. 1F), where the long and healthy mitochondria are better visualized than Figs. 1D & 1E with sharper boundaries. Accurate 3D imaging of mitochondria is essential to quantify mitochondrial volume. Fig. S2(A) shows the 3D renderings of the healthy mitochondria from the same volume as Fig. 1F. Fig. 1H shows an XZ cross-section of a mitochondrion captured using confocal microscopy, indicating that the hollow structure of the mitochondrial outer-membrane cannot be clearly resolved axially by confocal imaging. In the SMLM XZ cross-section (Fig. 1I), we observed that mitochondria interior regions had lower intensity than the boundaries due to their natural hollow structures, which are not evident in the confocal microscopy images (Fig. 1H).

SMLM’s high spatial resolution is particularly important in quantifying geometrical parameters in densely packed mitochondrial networks in cells. Fig. 1J is a confocal microscopy image of several densely packed mitochondria, where it is challenging to segment individual mitochondria due to poorly resolved mitochondrial boundaries. In contrast, SMLM resolved the boundaries between individual mitochondria (Fig. 1K,) setting the stage for statistically analyzing mitochondrial geometrical parameters. To further demonstrate the improved axial resolution in 3D SMLM, we show the depth profiles of two mitochondria stacked on top of each other in confocal microscopy (Fig. 1L) and SMLM (Fig. 1M) from the locations respectively highlighted in Fig. 1J and Fig. 1K. The depth profile from the confocal microscopy image suggests that is it nearly impossible to segment densely packed mitochondria due to diffraction-limited axial resolution. In contrast, The depth profile from the SMLM image shows three distinct peaks spaced ∼750 nm and 600 nm apart, corresponding to the mitochondrial membranes and matching published reports [51]. In Fig. S3, we compared more depth profiles of randomly selected individual mitochondria imaged by confocal microscopy and SMLM. Previous 3D electron microscopy studies of adherent HeLa cells showed that mitochondrial axial thickness typically ranges from 600 nm to 900 nm [51], which agrees with our SMLM measurements (Fig. S3A.) In comparison, our confocal microscopy measurement of mitochondrial axial thickness was over 1300 nm (Fig. S3B).

To characterize alterations in 3D SMLM images of HeLa cell mitochondria, we quantified the mitochondrial morphology and outer membrane protein distribution. We loaded the mitochondrial localization data into 3D 8-bit image files with 20 nm pixel size. We then isolated the mitochondrial signal from the background signal by applying an intensity threshold to each frame. The threshold was determined by

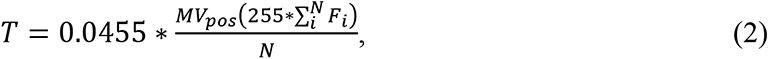

where *F* is the matrix of intensities in the frame; *i* is the pixel index; *N* is the total number of pixels in the frame; *MV*_*pos*_is the mean value of the positive pixels in the image; 0.0455 is an empirically determined factor; and 255 is the maximum pixel value in an 8-bit image. We formed 3D objects out of the thresholded image, filled the objects, filtered out objects of less than 0.05 μm^3^ in volume, then counted the total number of positive pixels to compute the total mitochondrial volume.

Figs. 2A&2B are results from the background removal via thresholding and the segmentation of overlapping mitochondria, respectively. Each mitochondrion is outlined in red in Fig. 2B, from which we calculated each mitochondrion volume. Figs. 2C&2D depict the skeletonization process of mitochondria, wherein each mitochondrion is collapsed to a single, one-dimensional line that traces the contour of its main axis. The resulting skeletons are shown in orange in Fig. 2C and follow the mitochondrial contours in all three dimensions, as shown in Fig. 2D. We used graph theory [52] to eliminate the branches of each mitochondrial skeleton (Fig. S4A). We designated each skeletal branchpoint and endpoint as a node connected to the other nodes by links (Fig. S4B), then identify the single longest 3D path connecting two nodes (Figs. S4C-S4F). We then treated this path as the central axis of the mitochondria and its length as the mitochondrial length.

**Fig. 2.**
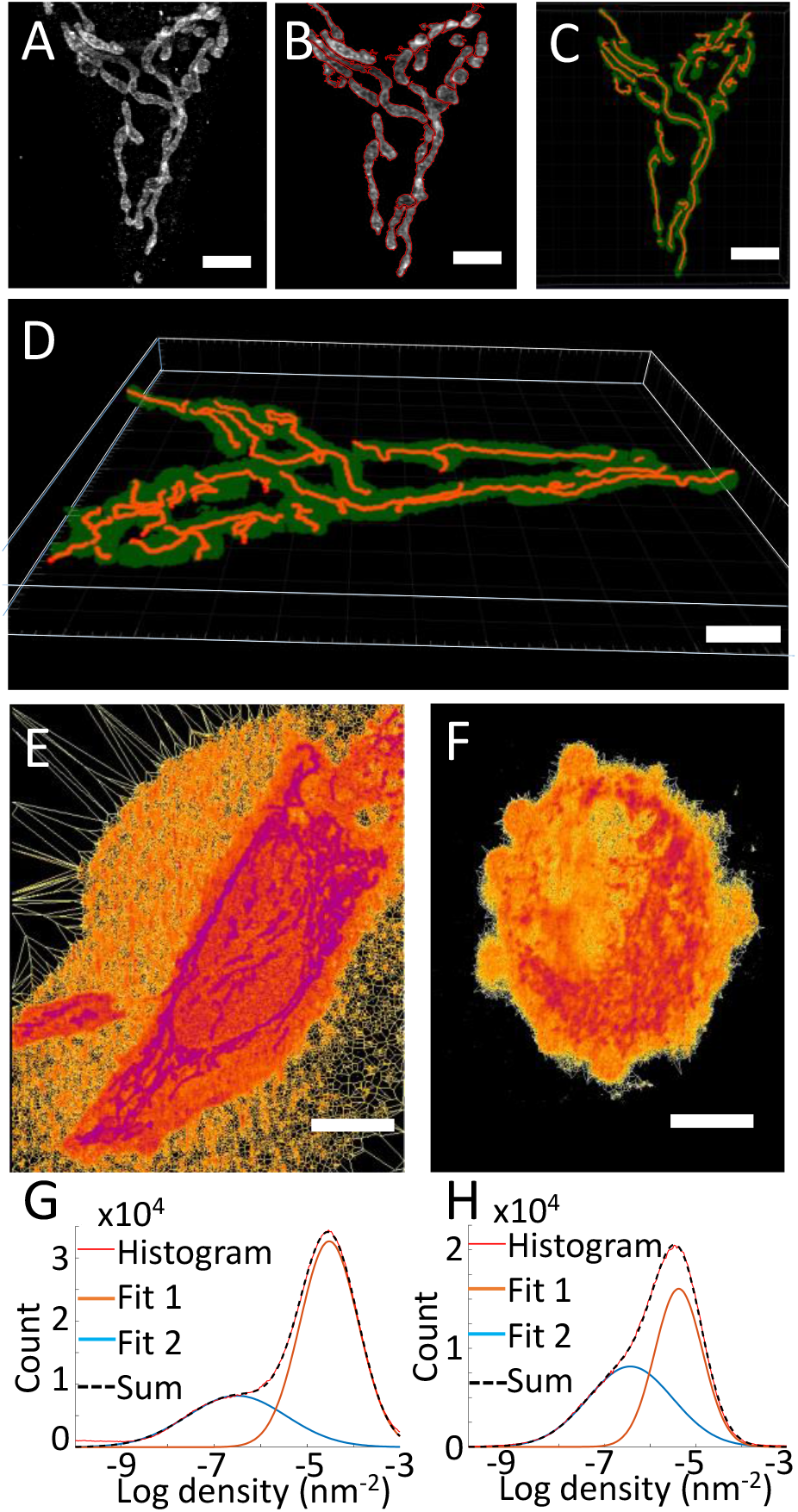
(A) Projection view of a mitochondria imaged by 3D SMLM. Scale bar: 3 μm; (B) The image from panel A with red outlines around each mitochondrion indicates segmentation results; (C) Skeletonized mitochondria from panel A (red) superimposed on a binarized image of the mitochondria (green); (D) Volumetric visualization of panel C; (E) Localization density-based analysis of a high-density (untreated) mitochondrial membrane structure. Scale bar: 10 μm; (F) Localization density-based analysis of a Low-density (STS-treated) mitochondrial membrane structure. Scale bar: 10 μm; (G) Histogram showing the log density of localizations of a compact membrane sample shown in panel E and two Gaussian functions fitted to the histogram contour. The blue line represents the estimated background density, the orange line represents the estimated true signal density, the black line is the sum of the blue and orange lines, and the red line is the original histogram contour; (H) Histogram showing the log density of localizations of a dissociated membrane sample (F) along with two Gaussian functions fitted to the histogram contour. The blue line represents the estimated background density, the orange line represents the estimated true signal density, the black line is the sum of the blue and orange lines, and the red line is the original histogram contour.

We measured the mean radius of the intersection between the plane perpendicular to the central mitochondrial axis at each point along the axis (Fig. S5A) and the mitochondrial outer membrane (Fig. S5B). We took the mean value of these measured radii as the mitochondrial width (Fig. S5C), then took the ratio of mitochondrial length to mitochondrial width to calculate the mitochondrial aspect ratio. We compared the quality of measuring the mitochondrial length and aspect ratio on 3D images and 2D projection images. We found that measurements from 2D projection images introduced significant variability in mitochondrial length (Fig. S6A-S6B), with a 25% lower average length and a standard deviation between 26% and 28% (Fig. S6C-S6D) compared with measurement from 3D images.

To analyze the protein localization density, we used 3D Voronoi tessellation [53], where borders are drawn around each localization corresponding to the equidistant points between the selected localization and its neighboring localizations. The local density in each molecule’s volume of influence is then determined by the inverse of the polygon volume, allowing us to measure changes in the density of TOM20 localizations. Fig. 2E shows a tessellated HeLa cell color-coded by localization density, in which smaller polygon volumes are purple and larger volumes are orange. Fig. 2F shows a tessellated HeLa cell with lower density.

To avoid the influence of factors such as antibody binding efficiency and blinking dynamics, we took the ratio of mitochondrial signal density to cytoplasmic signal density, which is associated with unbound TOM20 proteins [54] and nonspecific staining. Since both metrics increase with higher labeling density or binding efficiency, the ratio between them would remain constant. A lower ratio indicates dissociation of the mitochondrial membrane. We plotted a histogram of the log density, which contained two peaks, one associated with the mitochondrion-bounded TOM20 and one associated with unbound TOM20. We used least squares fit to fit two Gaussian distributions to each histogram, then subtracted the value of the low-density peak from the value of the high-density peak to get the ratio. This process is depicted in Fig. 2G, which shows a histogram of the log density of the image in Fig. 2E and the same histogram as approximated by two Gaussian functions fitted to it. These Gaussian functions are relatively distinct from each other, indicating a notable difference in densities between bound and unbound TOM20. In contrast, the two Gaussians in Fig. 2H, which shows the histogram of the log density of Fig. 2F, are more overlapped, indicating a smaller difference between the bound and unbound TOM20.

To validate our extraction of morphological parameters and protein density from mitochondria images, we tested our analyses on simulated ground-truth mitochondrial images. We simulated ground-truth 3D mitochondria structures using a worm-like chain model [55] with a lateral persistence length of 0.6 μm and axial persistence length of 4.8 μm. These parameters were extracted from our experimental mitochondrial images. To generate elongated (Fig. 3A), moderately elongated (Fig. 3B), and fragmented (Fig. 3C) mitochondria, we set the mitochondrial lengths to 4 μm, 1.5 μm, and 0.65 μm, respectively. To transform the worm-like chains from curved lines into objects with specified volumes, we convolved each defined point along the chain with an ellipsoid of radius

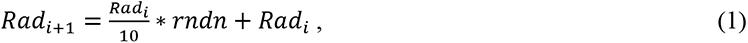

**Fig. 3.**
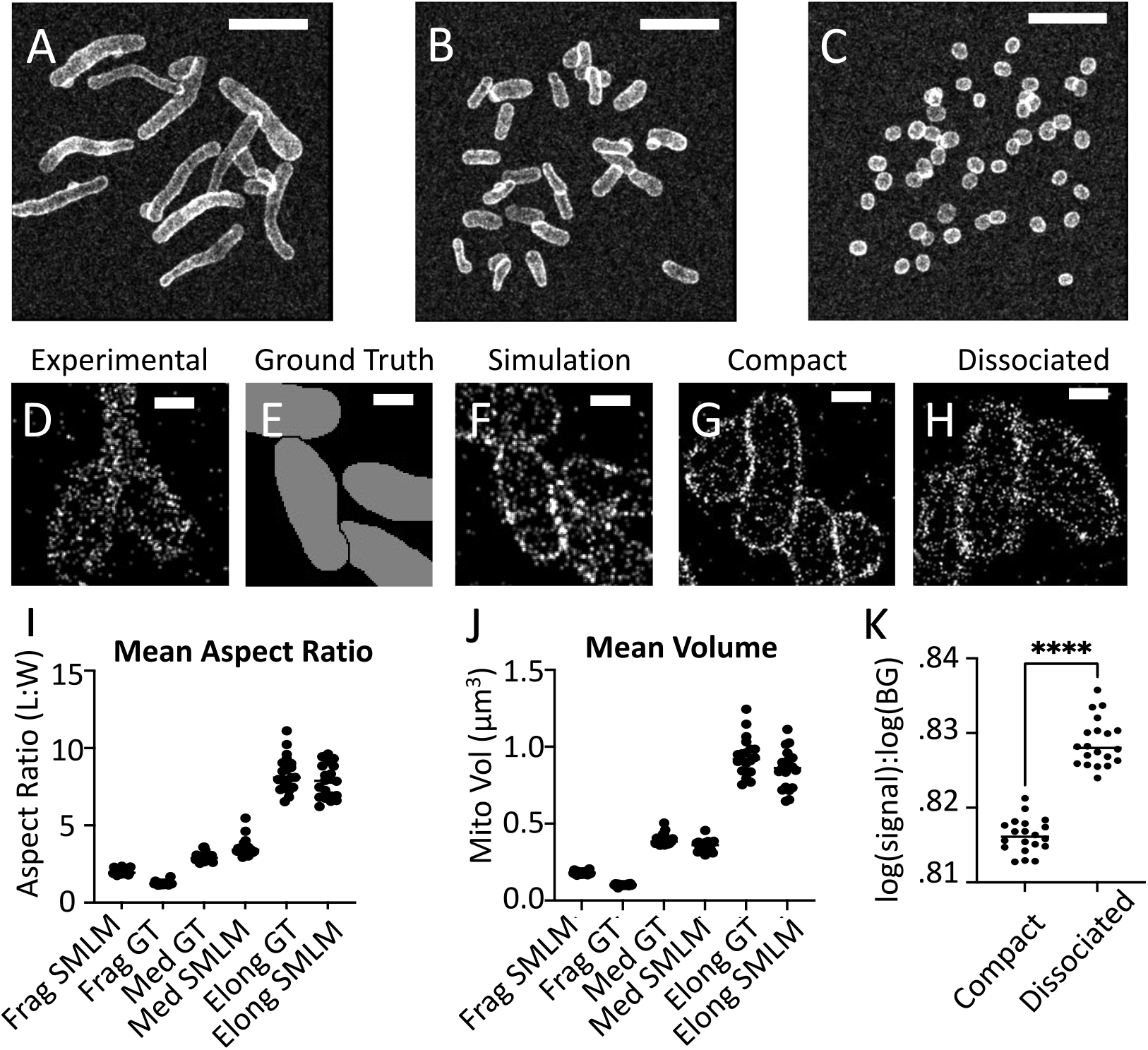
Simulated 3D SMLM images of (A) elongated (4.0 μm length), (B) moderately elongated (1.5 μm length), and (C) fragmented (0.65 μm length) mitochondria structures. Scale bar: 3 μm; (D) A 32-nm thick section of an experimental SMLM image of a healthy HeLa mitochondrion; (E) A 32-nm thick slice of a simulated ground truth mitochondrial shape. A 32-nm thick section of a simulated 3D SMLM mitochondrial image based on the generated shapes shown in panel E with (F) normal, (G) compact, and (H) disassociated outer membrane proteins. Scale bars: 500 nm; (I) Measured values of mitochondrial aspect ratio from simulated mitochondria with varying lengths (N=20); (J) Measured values of mitochondrial volume from simulated mitochondria with varying lengths (N=20); (K) Measured values of the ratio between the log signal density to log background density from simulated mitochondria with varying membrane dissociation levels (N=20).

where *i* is the position along the mitochondria; *Rad*_*i*_ is the mitochondrial radius at position *i* on the mitochondria; and *rnd*n is a random number generated from a Gaussian distribution with a mean of 0 and a standard deviation of 1. The initial radius of each ellipsoid was 0.325 μm in the lateral and axial directions. The lateral and axial radii at each point were determined independently. These parameters and relationships were generated empirically and iteratively by measuring parameters from experimental SMLM images of mitochondria. We generated 20-50 mitochondria per simulated image, depending on the extent of elongation. As shown in Fig. 3D, a 32-nm thick slice of an experimental 3D SMLM image, when the outer-membrane of two mitochondrion meet each other, they form an interface and do not overlap. To reflect this, if the chain of one mitochondrion overlapped with a previously established mitochondrion, we ceased its generation. This is shown in Fig. 3E, which shows that the simulated mitochondria do not overlap but rather meet each other then stop. These interfaces become important in recognizing mitochondrial boundaries during mitochondrial segmentation.

After simulating the ground-truth mitochondria, we simulated fluorescence emission and the imaging process using parameters from literature and experimental observation. We split the ground-truth volumes into 16×16×16 nm pixels. If a pixel contained the outer-mitochondrial membrane in a designated TOM20 cluster region, we gave it a 13% chance of containing at least one labeled TOM20 protein. If the pixel was located within the mitochondria, we gave it a 0.66% chance of containing at least one labeled protein. If the pixel was in the cytoplasm, we gave it a 0.125% chance of containing at least one labeled protein. These percentages were extrapolated from labeling densities reported by Wurm *et al.* [56] and from the observed ratios of label density in the cytoplasm and within the mitochondria in our SMLM images. This rule resulted in each simulated image containing roughly 150,000 labeled TOM20 proteins. Using a uniform distribution, we randomly translated these fluorophores within 17.5 nm of the ground truth location to reflect the size of the primary and secondary antibody probes [56, 57]. Based on observed localization numbers in our SMLM with similar size and mitochondrial density, we randomly generated 380,000 localizations from these 150,000 labeled proteins using an exponential distribution for the number of blinkings per localization [58]. We then applied lateral and axial uncertainties to each localization based on the observed parameters from maximum likelihood estimation (MLE) fitting in our experimental data. As a result, we could simulate 3D SMLM images (Fig. 3F) from our ground truth data.

To simulate outer-mitochondrial membrane dissociation, we applied an additional normal distribution to each ground-truth location with a sigma between 0 (Fig. 3G) and 35 nm (Fig. 3H). Fig. S7 contains a flowchart outlining the generation of simulated SMLM mitochondria images. We performed a Fourier Ring Correlation (FRC) analysis [59] with a resolution cutoff of 1/7 [60] to characterize the lateral resolution of the 2D projections of the SMLM images. We verified that the simulation conditions resulted in a lateral resolution of 65 nm, matching the FRC analysis of our experimental data (Fig. S8).

Next, we tested our geometrical and density analyses on the simulated 3D SMLM mitochondrial images. We found that the estimated aspect ratios matched well with the ground truth and, as expected, increased with more elongated mitochondria (Fig. 3I). We also found that the estimated volumes of the simulated mitochondria matched well with the ground truth (Fig. 3J). However, there was a systematically enlarged volume estimation compared to the ground truth. This bias is attributable to the lateral and axial localization uncertainties blurring the mitochondrial outer membrane and the primary and secondary antibody sizes [58]. However, it does not impede comparison between similar SMLM images since the bias universally affects all SMLM images. To validate our localization density analysis, we simulated one dataset with compact membrane proteins and one dataset with dissociated membrane proteins and measured their TOM20 densities using the method shown in Fig. 2. We found that simulated dissociated membranes had significantly lower signal density than the simulated compact membrane, as shown in Fig. 3k.

Figure 4 shows the validation of our geometrical analysis of mitochondrial fragmentation in HeLa cells treated with OA. OA treatment inhibits ATP synthase activity and leads to mitochondrial fragmentation and loss of MMP [61]. Fig. S9A shows our validation of MMP loss in OA-treated HeLa cells using JC-1 labeling. The measured MMP dropped significantly below the MMP of the untreated cells (Fig. 1C). Fig. 4A shows the 2D projection SMLM image of an OA-treated HeLa cell, and Fig. 4B shows the magnified view of the highlighted area in Fig. 4A. They both show fragmented mitochondria with high resolution. Confocal microscopy images of OA-treated HeLa cells also show a clear difference in mitochondrial morphology between the OA-treated and untreated HeLa cells (Figs. 4C&4D). The untreated cells exhibited relatively elongated mitochondria (Fig. 1D), and OA-treated cells showed fragmented mitochondria (Figs. 4B&4C). We also show a 3D visualization of one SMLM-imaged OA-treated cell (Fig. S2B). While confocal microscopy roughly showed the morphological features of the mitochondria, the individual mitochondrial hollow outer-membrane structures are unresolvable due to diffraction-limited axial resolution. We compare the cross sections of SMLM and confocal microscopy imaged OA-treated mitochondria in Figs. 4E&4F, which are extracted from the locations highlighted in Figs. 4B&4D, respectively. These cross-sectional views further validated that confocal microscopy has limited ability to resolve individual mitochondria fully and cannot accurately quantify their morphological parameters. Moreover, we found that the mitochondrial aspect ratio (Fig. 4G) and the total mitochondrial volume (Fig. 4H) became significantly lower in OA-treated cells.

**Fig. 4.**
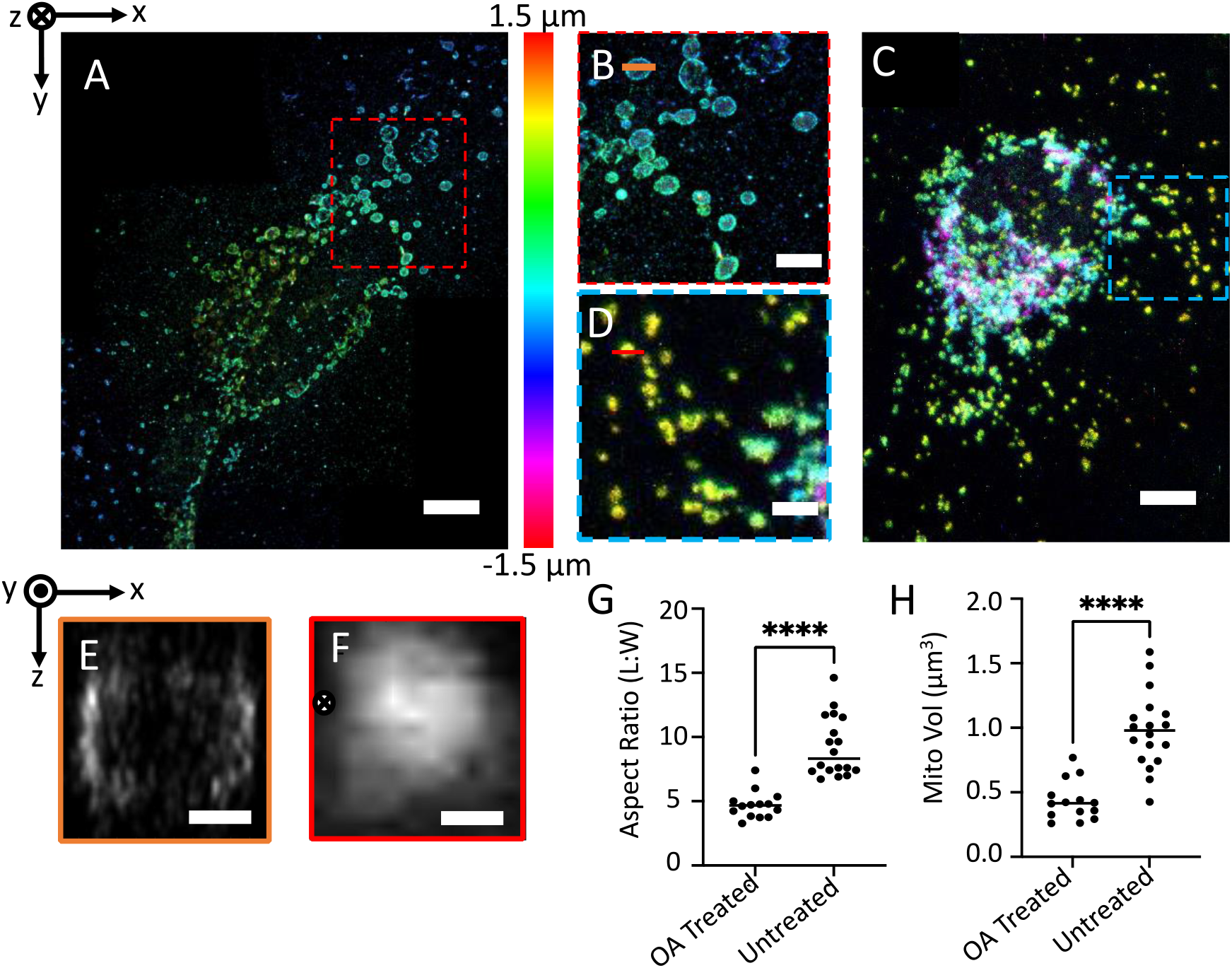
(A) 2D projection of a 3D SMLM image of OA-treated HeLa cell. Scale bar: 5 μm; (B) Magnified view of the area highlighted in panel A. Scale bar: 2.5 μm; (C) 2D projection of a 3D confocal microscopy image of OA-treated HeLa cell. Scale bar: 5 μm; (D) Magnified view of the area highlighted in panel C. Scale bar: 2.5 μm; (E) Cross-section of an SMLM imaged mitochondrion from the position highlighted in panel B. Scale bar: 250 nm; (F) Cross-section of a confocal microscopy imaged mitochondrion from the position highlighted in panel D. Scale bar: 250 nm; (G) Comparing mitochondrial aspect ratio between OA-treated and untreated groups from 3D SMLM (N=14 and N=18 for OA-treated and untreated groups respectively); (H) Comparing mitochondrial volume between OA-treated and untreated groups from 3D SMLM (N= 14 and 18 for OA-treated and untreated respectively).

Figure 5 shows the validation of our quantification of mitochondrial outer membrane decompaction in normal and STS-induced pre-apoptotic HeLa cells. Fig. 5A shows a whole STS-treated cell imaged by SMLM, and Fig. 5B shows a magnified view of the area highlighted in Fig. 5A. Both images show a more blurred mitochondrial outer membrane compared to untreated cells (Fig. 1K), indicating possible dissociation of the TOM20 proteins from the mitochondrial membrane. FRC analysis showed a comparable resolution between SMLM imaging of treated and untreated cells. In contrast, confocal microscopy images (Figs. 5C&5D) of STS-treated HeLa cell mitochondria did not exhibit significant changes in the compactness of the outer membrane. The 2D projections appeared similar to the confocal images of untreated cells (Fig. 1D&1E). This is likely due to the low resolution of the confocal images, which made them insensitive to changes in local protein density. Also, while the super-resolution cross-section of the STS-treated mitochondria still resolved the hollow structure (Fig. 5E), the confocal cross-sections did not (Fig. 5F). Using Voronoi tessellation, we found that the proteins were significantly denser in the untreated cells than in the STS-treated cells (Fig. 5G), suggesting membrane dissociation in the early stages of apoptosis. Our SMLM observation agrees with literature reports [40, 45]. Cells treated with STS also experienced higher MMP than untreated cells, as indicated by JC-1 fluorescence in the 590 nm range (Fig. S9B), agreeing with prior studies [62].

**Fig. 5.**
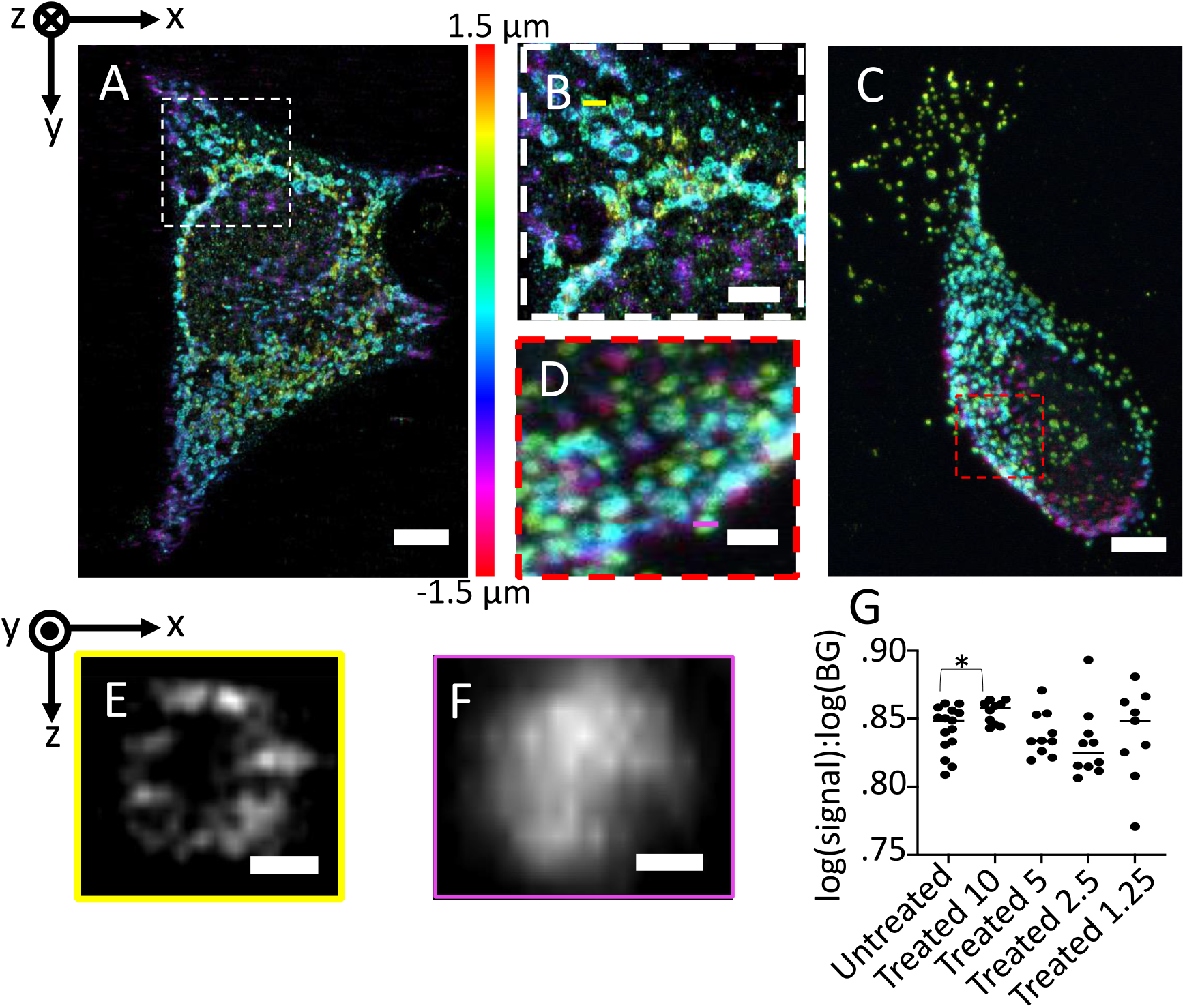
(A) 2D projection of a 3D SMLM image of STS-treated HeLa cell. Scale bar: 5 μm; (B) Magnified view of the area highlighted in panel A. Scale bar: 2.5 μm; (C) 2D projection of a 3D confocal microscopy image of an STS-treated HeLa cell. Scale bar: 5 μm; (D) Magnified view of the area highlighted in panel C. Scale bar: 2.5 μm; (E) Cross-section of an SMLM imaged mitochondrion from the position highlighted in panel B. Scale bar: 300 nm; (F) Cross-section of a confocal microscopy imaged mitochondrion from the position highlighted in panel D. Scale bar: 300 nm; (G) Comparison of membrane protein disassociation between untreated cells and cells treated with different concentrations of STS (N =15 in the untreated group, N=10 in the 10, 5, and 2.5 μM STS-treated groups, and N= 9 in the 1.25-μM STS-treated group).

### Discussion and conclusion

In this work, we developed methodologies to analyze the geometrical parameters of individual mitochondria, including volume and aspect ratio, for 3D SMLM. Using simulated ground-truth images, we validated the ability to detect alterations in individual mitochondrial morphology and outer mitochondrial membrane protein distribution. We further correlated these changes with chemically-induced perturbations in mitochondrial function using OA and STS. We found that mitochondria aspect ratio and volume decreased with OA treatment and that STS-treated cells had a lower mitochondrial outer membrane density than the untreated cells.

This validated method for evaluating individual mitochondrial health and function can identify conditions such as mitochondrial elongation, pointing to greater metabolic demands and mitochondrial damage from sources such as oxidative stress. It can also identify mitochondrial fragmentation, potentially correlated with a decreased metabolic demand or ability, or cells in a quiescent or apoptotic state [21, 25]. Finally, it can identify the density of mitochondrial outer-membrane signal, another metric of impending apoptosis and outer-membrane damage [45]. Applying our method may help to evaluate the role and extent of mitochondrial dysfunctions in various metabolic diseases.

Our method is unique among existing mitochondrial morphology analysis tools because it exploits the fine features only resolvable by super-resolution microscopy. Namely, it can quantify mitochondria individually and quantify the distribution of proteins on the mitochondrial membrane. In addition, other groups have shown that it is possible to resolve mitochondrial cristae using super-resolution microscopy [63-65]. Our analytical method could be expanded in the future by quantifying parameters such as cristae surface area, which gives a more direct measure of mitochondrial metabolism since mitochondrial respiration takes place on the cristae [66]. The current weakness of our method comes from manual segmentation. We currently rely on manual segmentation of individual mitochondria; hence analyzing a large number of cells is highly labor-intensive and time-consuming. In the future, individual mitochondria segmentation can greatly benefit from machine-learning-based algorithms to automatically segment overlapping mitochondria and significantly improve the analysis throughput.

## SUPPLEMENTARY MATERIALS

**Fig. S1:**
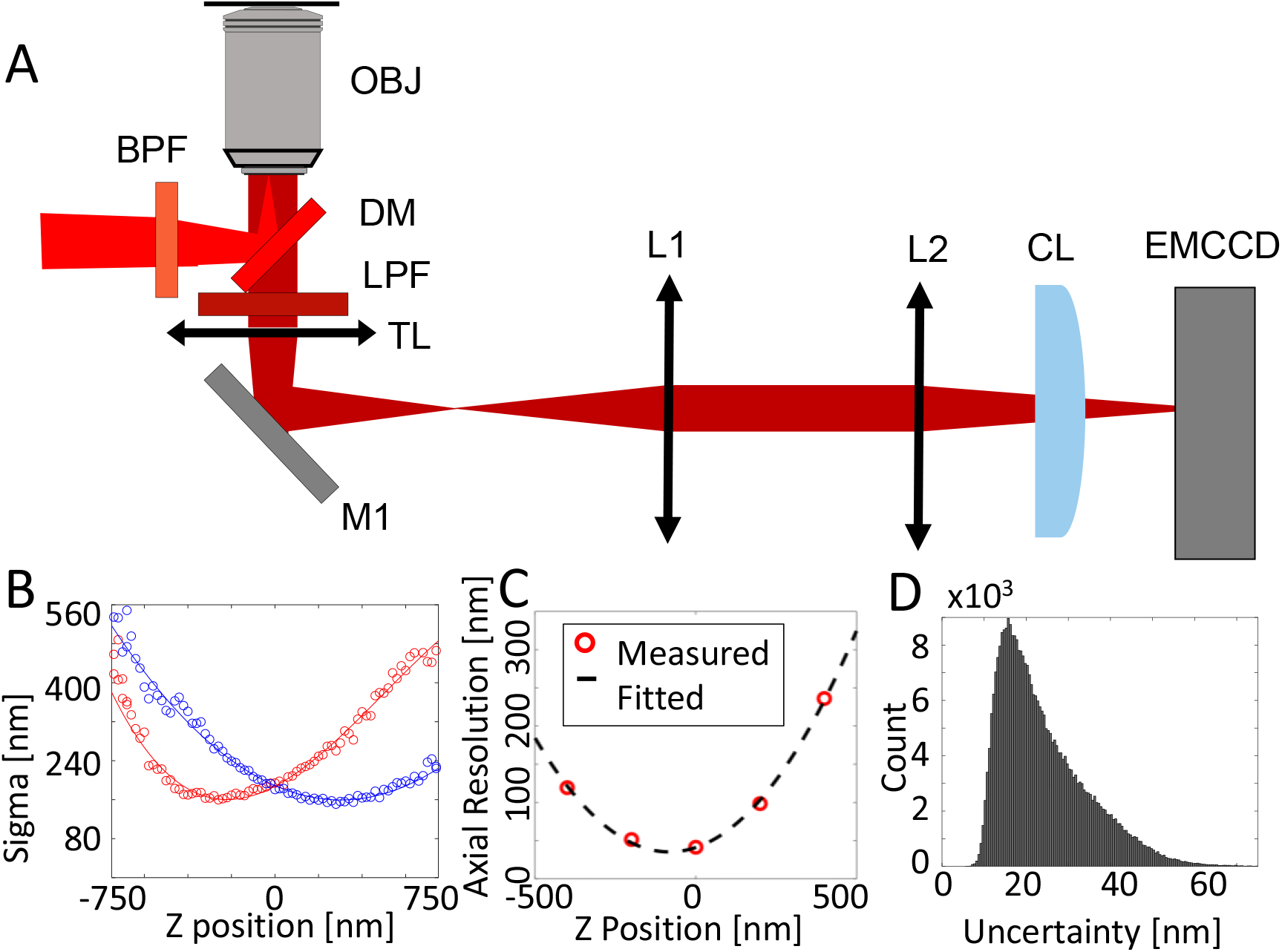
(A) Schematic of the 3D SMLM system. OBJ: objective lens; BPF: band-pass filter; DM: dichroic mirror; LPF: long-pass filter; TL: tube lens; M: mirror; L: lens; CL: cylindrical lens; (B) Calibration curve for axial localization showing the estimated sigma of the PSF in the x- and y-directions at different depths; (C) Measured axial resolution as a function of the axial position of a nanosphere with 2,000 photon count; (D) Histogram of estimated sample localization uncertainty (median uncertainty: 17 nm).

**Fig. S2:**
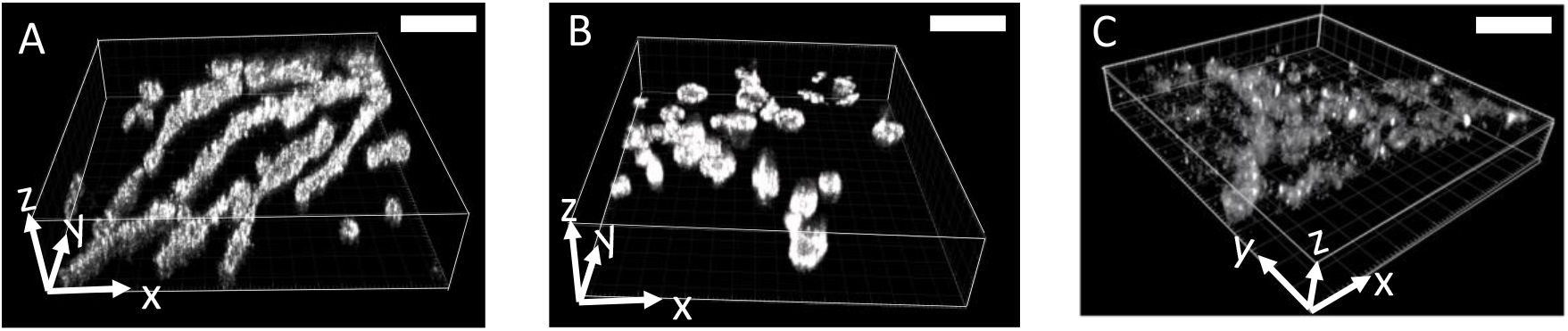
Volumetric visualization of SMLM imaged mitochondria from (A) Normeal HeLa cells; (B) STS-treated HeLa cell; (C) OA-treated HeLa cells. Scale bar: 3 μm. A

**Fig. S3:**
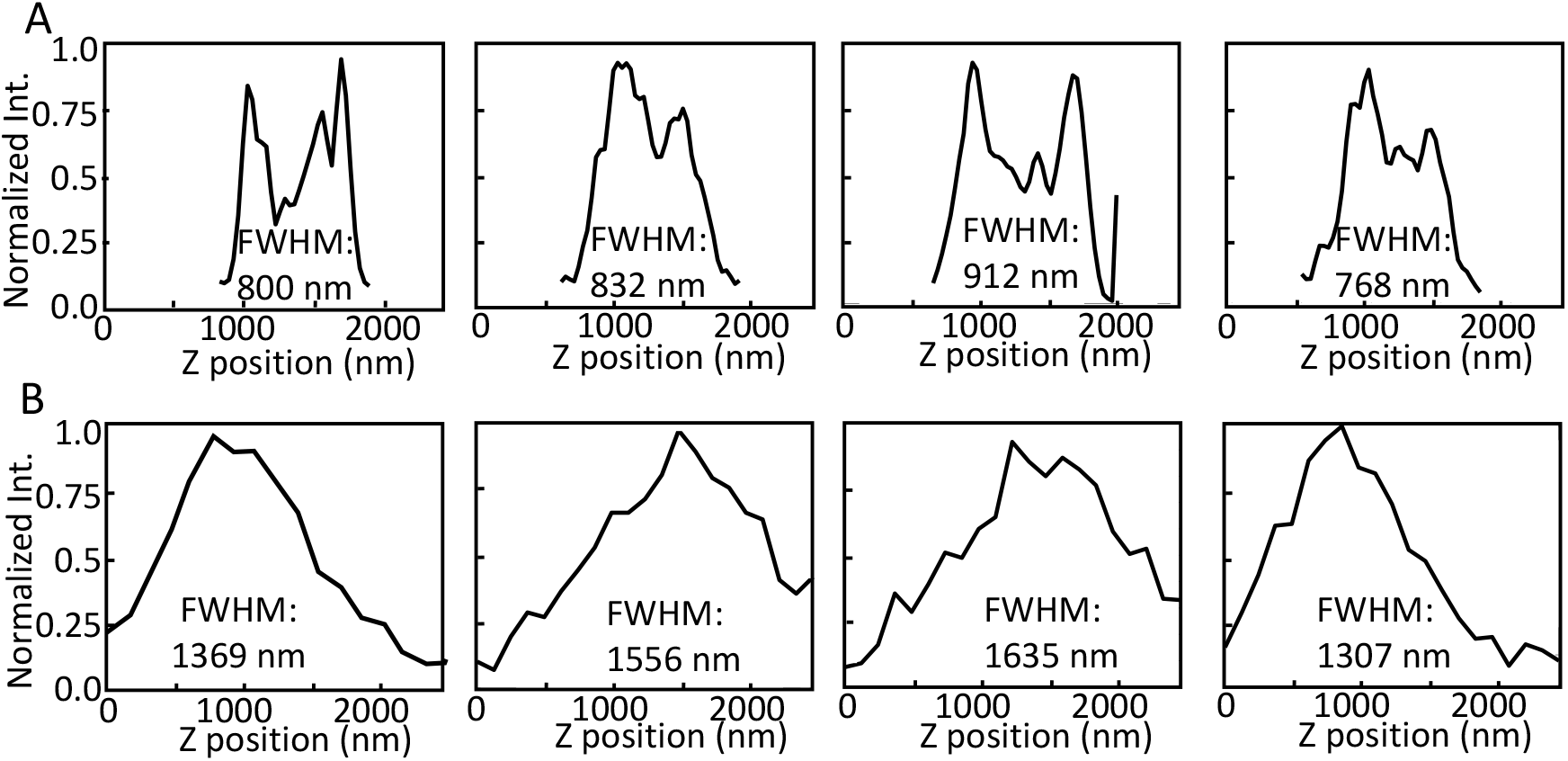
**(A**) Representative depth profiles of mitochondria imaged by 3D SMLM; (B) Representative depth profiles of mitochondria imaged by 3D confocal microscopy.

**Fig. S4:**
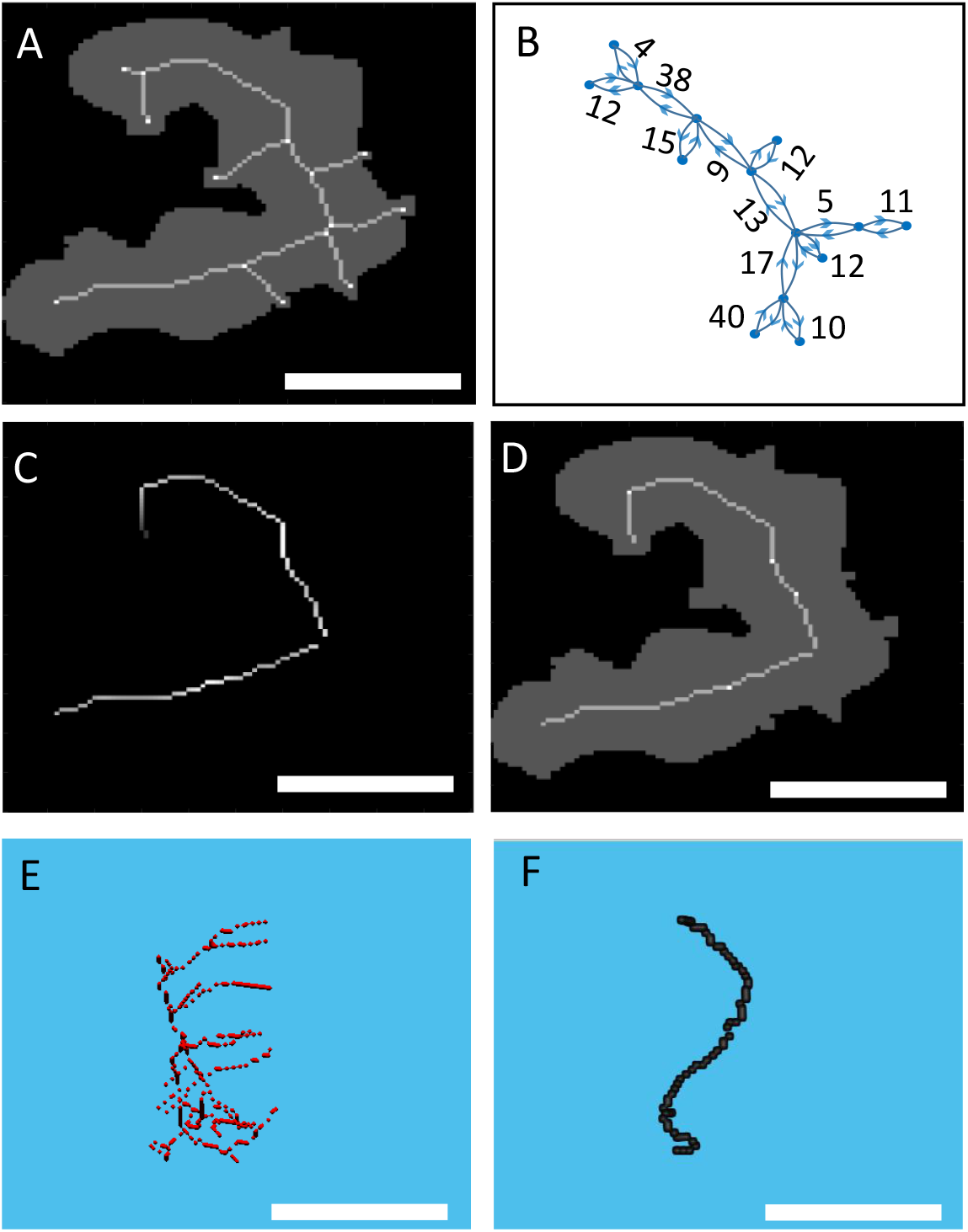
(A) Skeletonized simulated 2D mitochondria with branches; (B) Decomposition of the skeleton into nodes and links, annotated with distances between each node; (C) The isolated single main path of the mitochondria after elimination of minor branches. (D) the Main branch of the mitochondria superimposed upon the original structure. (E) The skeleton of an experimental 3D image of a mitochondrion. (F) The main path of the mitochondria from (E) with minor branches removed. Scale bars: 750 nm

**Fig. S5:**
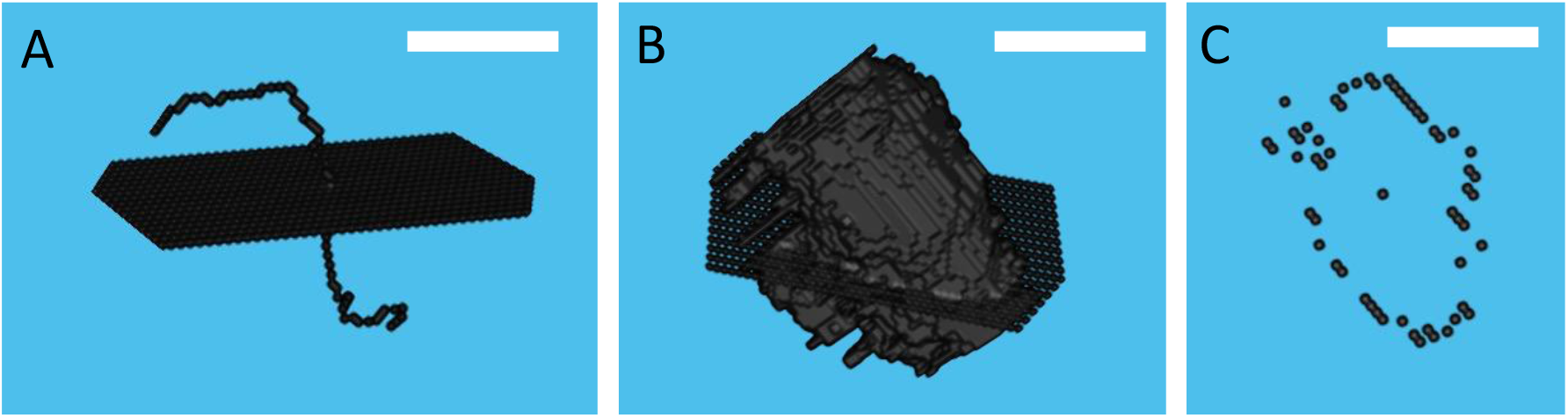
Procedure to measure mitochondrial width. (A) A plane is drawn perpendicular to the mitochondrial skeleton at each point; (B) The outer surface of the mitochondria overlaps the drawn plane from panel A; (C) The intersection points between the perpendicular plane and the mitochondrial outer surface, as well as the center point on the mitochondrial skeleton. Scale bars: 750 nm.

**Fig. S6:**
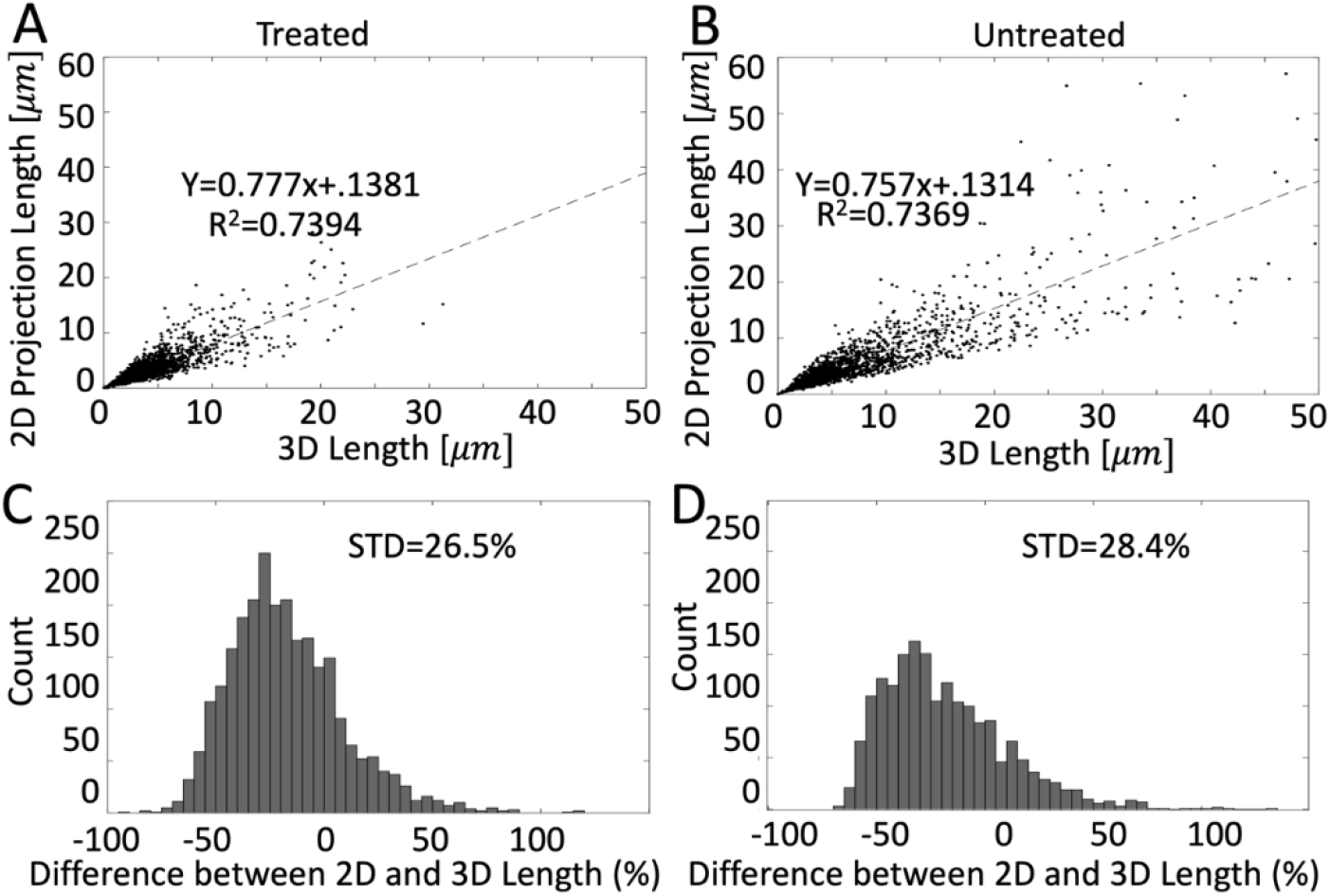
Comparing mitochondria length measured from 2D projection and 3D images in (A) oligomycin-treated and (B) untreated cells. The dashed lines represent the best linear fit. Histogram of the difference between mitochondria length measured from 2D projection and 3D images in (C) oligomycin-treated and (B) untreated cells.

**Fig. S7:**
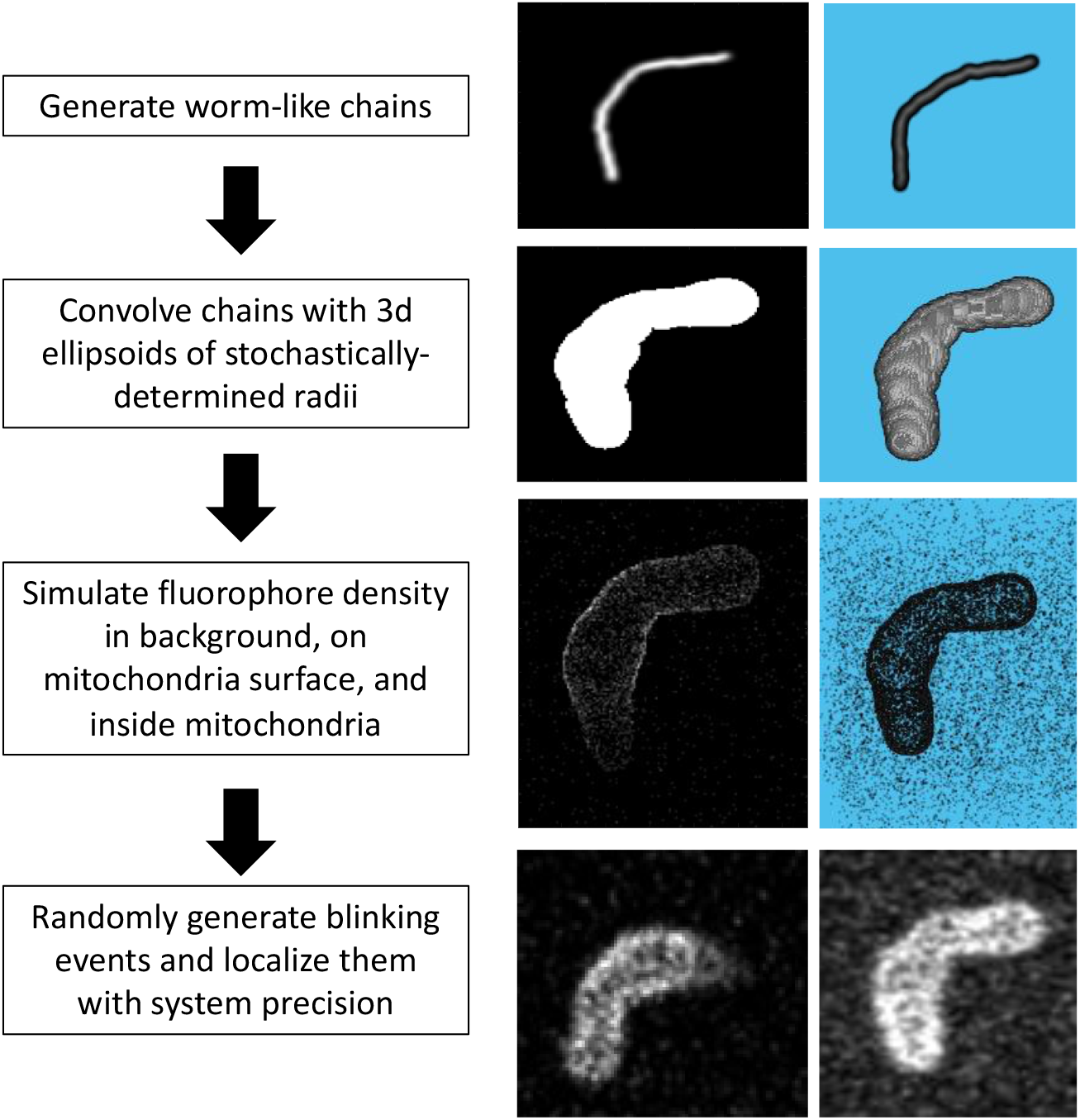
Flowchart (left column) describing the steps of generating simulated mitochondria, starting from a worm-like chain and ending with a simulated 3D SMLM image. Illustrations of a 100-nm slice of a mitochondrion (middle column) and a full 3D mitochondrion (right column) corresponding to each step in the flow chart.

**Fig. S8:**
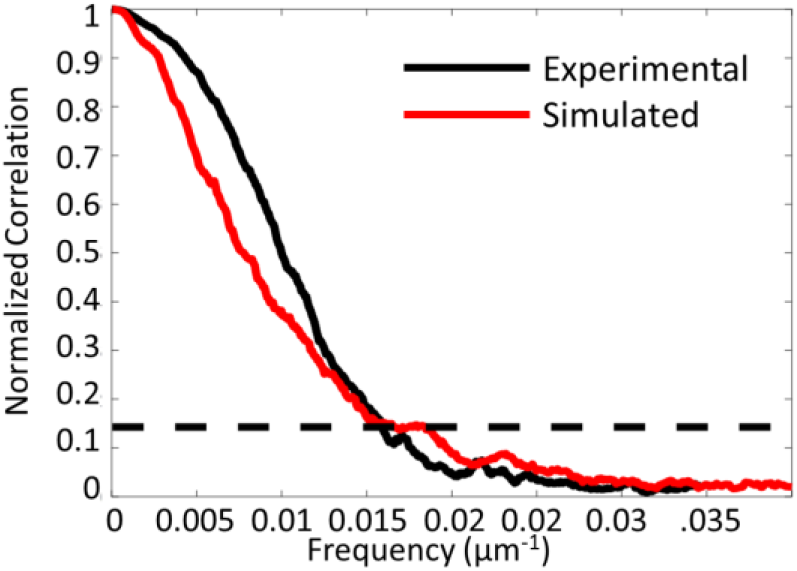
FRC results of experimental and simulated results with 380,000 localizations from 150,000 unique labels. The FRC resolution is 65 nm.

**Fig. S9:**
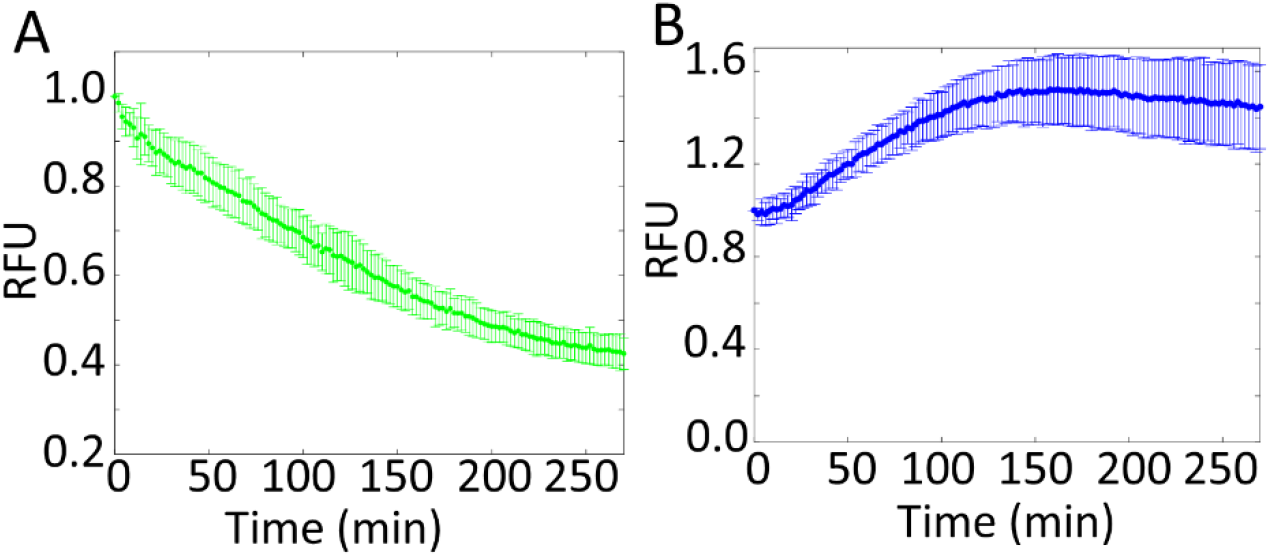
(A) Normalized RFU over time in HeLa cells labeled with JC-1 and treated with OA. (B) Normalized RFU over time in HeLa cells labeled with JC-1 and treated with 10-μM STS.

## Notes

### Competing Interest Statement

The authors have declared no competing interest.

